# Thoracic Aortic Three-Dimensional Geometry

**DOI:** 10.1101/2024.05.09.593413

**Authors:** Cameron Beeche, Marie-Joe Dib, Bingxin Zhao, Joe David Azzo, Hamed Tavolinejad, Hannah Maynard, Jeffrey Duda, James Gee, Oday Salman, Penn Medicine BioBank, Walter R. Witschey, Julio A. Chirinos

## Abstract

**Background:** Aortic structure impacts cardiovascular health through multiple mechanisms. Aortic structural degeneration occurs with aging, increasing left ventricular afterload and promoting increased arterial pulsatility and target organ damage. Despite the impact of aortic structure on cardiovascular health, three-dimensional (3D) aortic geometry has not been comprehensively characterized in large populations.

**Methods:** We segmented the complete thoracic aorta using a deep learning architecture and used morphological image operations to extract multiple aortic geometric phenotypes (AGPs, including diameter, length, curvature, and tortuosity) across various subsegments of the thoracic aorta. We deployed our segmentation approach on imaging scans from 54,241 participants in the UK Biobank and 8,456 participants in the Penn Medicine Biobank.

**Conclusion:** Our method provides a fully automated approach towards quantifying the three-dimensional structural parameters of the aorta. This approach expands the available phenotypes in two large representative biobanks and will allow large-scale studies to elucidate the biology and clinical consequences of aortic degeneration related to aging and disease states.

## 1. Introduction

The aorta is the largest conduit artery in the human body^1^. In addition to its conduit function, the aorta plays a key role in modulating pulsatile arterial hemodynamics, mediated by its cushioning function of the intermittent left ventricular ejection. Aortic structural parameters (including geometry and wall stiffness) have been shown to be key determinants of aortic hemodynamic function^1,2^. Despite its prominent age-associated changes and its key hemodynamic role, studies related to structural properties of aortic geometric phenotypes (AGPs) are lacking^3,4^. For example, previous cross-sectional studies among 210-250 patients undergoing aortic imaging demonstrated that the aorta elongates with age and that thoracic aortic length is greater among patients with acute aortic dissection.^3,4^ Moreover, other studies have been limited to two-dimensional cross-sectional geometric parameters of the aorta, neglecting important three-dimensional (3D) aspects of its geometry, including elongation, tortuosity/unfolding, and curvature, all of which influence aortic function.

Three-dimensional tomographic imaging (i.e. magnetic resonance imaging, computed tomography) has become a valuable resource for quantifying structural properties of the heart. Previous analyses of cardiac phenotypes acquired through the segmentation of CMR imaging data have developed a comprehensive atlas of cardiac structure and function in over 50,000 participants of the UK Biobank^5,6^; however, such analyses of three-dimensional aortic geometry have not been performed to date.

In this study, we present a deep learning approach to segment and comprehensively characterize the three-dimensional geometry of the thoracic aorta, and measure key phenotypes (diameter, length, curvature, tortuosity/unfolding) across various thoracic aortic subsegments. Our segmentation approach consists of **(1)** modality specific image segmentation, and **(2)** 3D mesh phenotype extraction. We deployed our segmentation approach on imaging data from 54,241 participants enrolled in the UK Biobank^7^ (UKB) and 8,456 individuals enrolled in the Penn Medicine Biobank^8^ (PMBB).

## 2. Methods

### Data Sources

The UKB is composed of >500,000 participating individuals aged 37-73 years at the time of recruitment, who underwent various questionnaires, physical measurements, biological sampling (blood and urine), and genome sequencing across 22 assessment centers in the UK^9^. A subset of participants were invited to complete an additional examination that included magnetic resonance imaging of the heart^7^. Data from a subset of individuals (n=54,241) enrolled in the UKB who underwent cardiac magnetic resonance imaging (CMR) were included for this analysis^7^ (**Table 1**).

The PMBB is composed of >250,000 consenting patients of the Penn Medicine health network^8^. Additionally, all participants medical records including imaging results are de-identified and linked to their identifier. We identified 8,456 patients enrolled who had thoracic computed tomography (CT) imaging data (**Table 1**).

### Aortic segmentation

We created a dataset of 233 axial steady-state free precision (SSFP) CMR images from the UKB to train and validate a deep learning segmentation algorithm to delineate the thoracic aorta. All CMR images were manually segmented by two trained medical doctors using in-house software. A variation of the U-Net convolutional neural network (CNN) segmentation architecture was used to segment the aorta^10^. The U-Net architecture is an encoder-decoder consisting of two-dimensional convolutional layers of increasing depth (shown in Fig. 2.). The encoder consisted of sequential convolutional and max-pooling layers, allowing the network to learn feature-representations across multiple spatial representations. The decoder consisted of an equivalent implementation with max-pooling layers being supplemented for up-sampling layers. Furthermore, all encoder feature-representations were concatenated onto their corresponding decoder feature-representations. The model was trained on 194 manually segmented CMR scans and validated on an independent subset of 39 CMR scans.

**Fig. 1.**
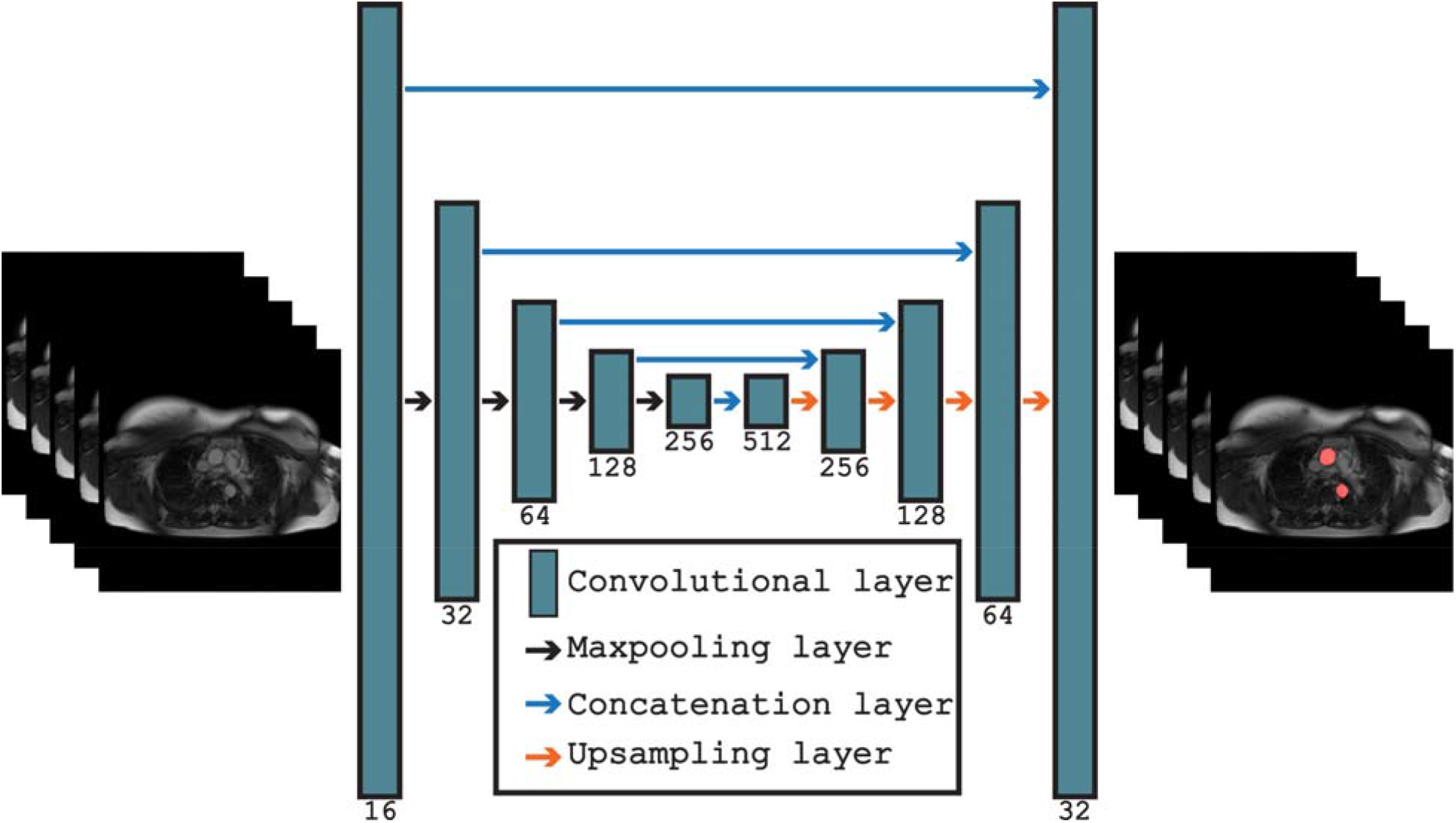
Overview of the U-Net segmentation architecture for performing segmentation of axial MRI images in the UK Biobank.

**Fig. 2.**
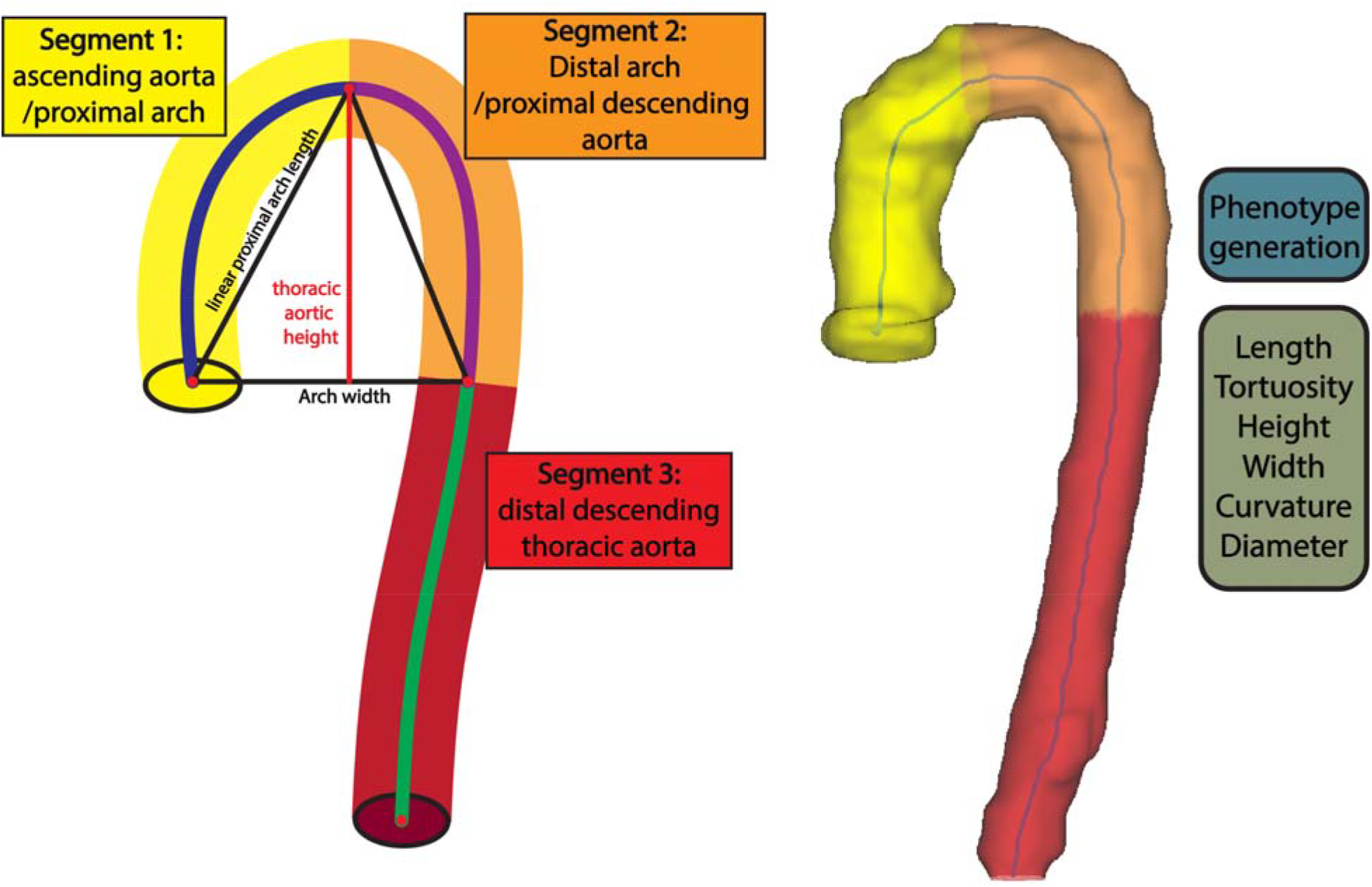
(A) Aortic mesh legend for each AGP region. (B) Visualization of three-dimensional aortic mesh.

The network was trained using the dice similarity coefficient (DSC) as the loss function and was optimized using the Adam optimizer with a learning rate of 0.001^11^. Training was performed for 20 epochs with a batch-size of 16. Image preprocessing consisted of zero-padding the 238×238 axial images to 240x240 followed by geometric and intensity augmentations being randomly performed on the training data, and Z-score normalization. Axial CMR image slices were segmented individually before being merged to form the three-dimensional aorta segmentation. The model achieved an average DSC of 0.934 (0.013) on the validation dataset. PMBB CT-scan data was segmented using the previously developed Totalsegmentator algorithm^12^. To extract the thoracic aorta region, the segmentation was cut at the T12 vertebral level.

### Three-dimensional aortic geometric phenotyping

The voxelized aorta segmentation was then post-processed using morphological erosion and dilation operations with a kernel size of 3 voxels to remove any holes in the segmentation. Mesh generation and centerline extraction were performed using the VMTK toolkit^13^. Iterative smoothing of the three-dimensional aortic mesh was performed using the Taubin algorithm^14^.

The centerline of the aortic mesh was then interpolated to have 100 points using B-spline interpolation. Identification of aortic subsegments was accomplished through exploiting properties of the vertical axis (z-axis) of the aorta’s centerline to identify: **(1)** the apex of the aorta, defined as the maximum centerline point on the vertical axis; **(2)** the centerline point on the descending aorta with smallest Euclidean distance on the vertical axis with the aortic root. The segment extending from the root to the apex of the aorta was identified as a single segment, hereby named ascending aorta/proximal arch. Similarly, the segment from the aortic apex to the point on the descending aorta vertically corresponding to the level of the aortic root was identified as a single segment, hereby named distal arch/proximal descending aorta. We note that these do not necessarily correspond to standard anatomical segmental definitions, since the latter are defined by the location of the major arch branches, which have variable relationships with the three-dimensional aortic apex^3^. Figure 2.a provides an aortic geometric phenotype key to understand each region, whereas Figure 2.b provides a three-dimensional mesh segmentation.

Centerline length was computed as the summation of distances between consecutive points. Minimal linear length was computed as the distance between the first and last point within each aortic segment. Tortuosity was computed as the ratio of centerline length to minimal linear length minus 1:

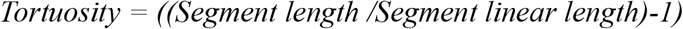

Aortic curvature was calculated using the VMTK toolkit^13^. The aortic radius was computed across every point on the centerline by calculating the nearest distance from the centerline to the aorta mesh and doubled to compute short-axis aortic diameter.

In addition to cross-sectional diameter, centerline length, tortuosity, and curvature, various key AGPs were computed from the overall aorta and from specific subsegments using the 3D-mesh. Given that the aortic arch and adjacent aortic segments are expected to be curved, standard curvature and tortuosity metrics do not intuitively characterize the expected aortic geometry. Therefore, to characterize thoracic aortic geometry, we used length, height and width of the segment from the aortic root to the corresponding vertical level in the descending thoracic aorta. We also computed the thoracic aortic unfolding index as follows:

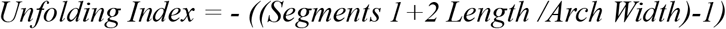

### Inferencing in the UKB and PMBB

We automatically segmented the entire thoracic aorta and derived 20 unique 3D-AGPs from 54,241 UKB participants and 8,456 PMBB participants (**Table 2**).

## 3. Discussion

This study leveraged aortic imaging data from two large independent cohorts, the UKB and the PMBB, to comprehensively characterize three-dimensional structural parameters of the thoracic aorta. We developed, for the first time, a phenotype extraction protocol consisting of **(1)** a deep learning segmentation architecture that accurately segments the thoracic aorta, and **(2)** morphological operations that operate directly on three-dimensional mesh representations to compute aortic length, diameter, tortuosity, curvature and arch measurements, as well as identify subsegments of the aorta. Our UKB segmentation model generated a DSC performance of 0.934, whereas the previously developed Totalsegmentator generated a DSC of 0.981^12^. We then successfully deployed our segmentation approach on 54,241 UKB imaging participants and 8,456 PMBB participants (**Table 2**). Our method provides reliable and quantitative phenotyping of three-dimensional aortic phenotypes. The two-stage segmentation approach allows for AGPs to be computed for any aortic segmentation map regardless of its initial modality. This method provides a fully automated approach towards quantifying the three-dimensional structural parameters of the aorta and expands the available phenotypes in two large representative biobanks. This will allow large-scale studies to elucidate the genetics, biologic mechanisms and clinical consequences of aortic degeneration related to aging and disease states in humans.

Our method is limited by the image resolution of our tomographic imaging data. Specifically, we were unable to identify the brachiocephalic and left subclavian artery to biologically delineate the aortic arch; however, we were able to provide geometric alternatives that approximate the anatomical regions in question. We note that the aortic subsegments do not necessarily correspond to standard anatomical segmental definitions or aortic subsegments with different embryological origins but were rather derived based on objective unequivocal landmarks in the three-dimensionally segmented thoracic aorta. In particular, the segment extending from the root to the apex of the aorta (segment 1 in **Fig. 2.a**), comprising the proximal transverse aortic arch, was identified as a single segment. Similarly, the segment from the apex to the plane intersecting the aortic root (segment 2 in **Fig. 2.c**) was identified as a single segment, whereas the sum of the two segments above were used to calculate the height and width of segments 1+2. Given that this height computation incorporates the ascending aorta and the proximal descending aorta, we note that these height and width indices from our study are not directly comparable to previous studies that measured aortic arch height and width starting from an arbitrarily defined plane intersecting some point of the middle of the anatomic ascending and descending aorta^15^.

In conclusion, we present a novel segmentation approach that accurately quantifies the three-dimensional geometry of the thoracic aorta, key subsegments, and provides comprehensive phenotyping of key 3D aortic geometric properties. We leverage our segmentation approach to perform inferencing on all participants in the UK Biobank and Penn Medicine Biobank with available imaging data. Future work should investigate the prognostic value as well as the biological foundation of AGPs.

## Acknowledgements

We acknowledge the Penn Medicine BioBank (PMBB) for providing data and thank the patient-participants of Penn Medicine who consented to participate in this research program. We would also like to thank the Penn Medicine BioBank team and Regeneron Genetics Center for providing genetic variant data for analysis. The PMBB is approved under IRB protocol# 813913 and supported by Perelman School of Medicine at University of Pennsylvania, a gift from the Smilow family, and the National Center for Advancing Translational Sciences of the National Institutes of Health under CTSA award number UL1TR001878.

## Supplemental Note

Penn Medicine BioBank Banner Author List and Contribution Statements

## PMBB Leadership Team

Daniel J. Rader, M.D., Marylyn D. Ritchie, Ph.D.

### Contribution

All authors contributed to securing funding, study design and oversight. All authors reviewed the final version of the manuscript.

## Patient Recruitment and Regulatory Oversight

JoEllen Weaver, Nawar Naseer, Ph.D., M.P.H., Giorgio Sirugo, M.D., P.h.D., Afiya Poindexter, Yi-An Ko, Ph.D., Kyle P. Nerz

### Contributions

JW manages patient recruitment and regulatory oversight of study. NN manages participant engagement, assists with regulatory oversight, and researcher access. GS assists with researcher access. AP, YK, KPN perform recruitment and enrollment of study participants.

## Lab Operations

JoEllen Weaver, Meghan Livingstone, Fred Vadivieso, Stephanie DerOhannessian, Teo Tran, Julia Stephanowski, Salma Santos, Ned Haubein, P.h.D., Joseph Dunn

### Contribution

JW, ML, FV, SD conduct oversight of lab operations. ML, FV, AK, SD, TT, JS, SS perform sample processing. NH, JD are responsible for sample tracking and the laboratory information management system.

## Clinical Informatics

Anurag Verma, Ph.D., Colleen Morse Kripke, M.S. DPT, MSA, Marjorie Risman, M.S., Renae Judy, B.S., Colin Wollack, M.S.

### Contribution

All authors contributed to the development and validation of clinical phenotypes used to identify study subjects and (when applicable) controls.

## Genome Informatics

Anurag Verma Ph.D., Shefali S. Verma, Ph.D., Scott Damrauer, M.D., Yuki Bradford, M.S., Scott Dudek, M.S., Theodore Drivas, M.D., Ph.D.

### Contribution

AV, SSV, and SD are responsible for the analysis, design, and infrastructure needed to quality control genotype and exome data. YB performs the analysis. TD and AV provides variant and gene annotations and their functional interpretation of variants.

## Statement of Ethics

This research has been conducted using the UK Biobank Resource under Application Number 81032. The PMBB is approved under IRB protocol# 813913.

## Author Contributions

C.B: Writing, methodology, and data analysis

M.J.D: Writing and data analysis

B.Z: Review and interpretation

J.D.A: Data preparation

H.T: Writing and review

H.M: Data preparation

J.D: Data analysis

J.G: Data acquisition and funding

O.S: Data preparation

PMBB: Data acquisition and funding

W.R.W: Data acquisition and funding

J.A.C: Conceptualization, writing, and funding

## Data Availability Statement

UK Biobank data is available to researchers following UKB application approval and IRB approval. Researchers can apply at https://www.ukbiobank.ac.uk/. Access to Penn Medicine BioBank data is provided to investigators at the University of Pennsylvania. External access can be provided through collaboration with an investigator at the university. Access to any code used in this analysis will be made available on reasonable request.

