## Supplementary material for "Thoracic Aortic Three-Dimensional Geometry": Tables

**Table 1.** Sample Characteristics in the UK Biobank and the Penn Medicine Biobank.

|  | UK Biobank (n=54,241) |  | Penn Medicine Biobank (n=8,456) |
| --- | --- | --- | --- |
| **Variable** |  |  |  |
| **Age** | 65.36 (±7.73) |  | 60.1 (±17.7) |
| **Male sex** | 26,002 (47.93%) |  | 4,516 (53.40%) |
| **Height (cm)** | 169.67 (±9.10) |  | 171.06 (±10.32) |
| **Weight (kg)** | 76.87 (±14.83) |  | 86.21 (±22.37) |
| **BMI (kg/m2)** | 26.61 (±4.21) |  | 29.42 (±6.98) |
| **Smoker** | 3,376 (6.22%) |  | 3,852 (38.5%) |
| **Systolic blood pressure (mmHg)** | 136.81 (±18.67) |  | 128.65 (±18.99) |
| **Diastolic blood pressure (mmHg)** | 81.43 (±10.41) |  | 76.63 (±11.68) |

**Table 2.** Descriptive statistics for aortic geometric phenotypes in the UK Biobank and Penn Medicine Biobank.

|  | **UK Biobank (n=54,241)** | | | |  | **Penn Medicine Biobank (n=8,456)** | | | |
| --- | --- | --- | --- | --- | --- | --- | --- | --- | --- |
| **Phenotype** | Mean | Median | Lower | Upper |  | Mean | Median | Lower | Upper |
| **Diameter** |  |  |  |  |  |  |  |  |  |
| Proximal arch diameter | 2.26 (±0.30) | 2.26 | 1.67 | 2.85 |  | 2.71 (±0.39) | 2.71 | 1.95 | 3.46 |
| Segment 1+2 diameter | 2.21 (±0.24) | 2.21 | 1.74 | 2.68 |  | 2.56 (±0.35) | 2.56 | 1.87 | 3.24 |
| Segment 2+3 diameter | 2.07 (±0.21) | 2.06 | 1.66 | 2.49 |  | 2.30 (±0.35) | 2.30 | 1.61 | 2.98 |
| Thoracic aorta diameter | 2.12 (±0.21) | 2.11 | 1.71 | 2.53 |  | 2.42 (±0.33) | 2.43 | 1.78 | 3.07 |
| **Length** |  |  |  |  |  |  |  |  |  |
| Proximal arch length | 8.52 (±1.90) | 8.44 | 4.80 | 12.24 |  | 10.54 (±2.39) | 10.49 | 5.86 | 15.21 |
| Segment 1+2 length | 17.13 (±2.89) | 17.06 | 11.47 | 22.79 |  | 21.07 (±3.91) | 21.05 | 13.40 | 28.74 |
| Segment 2+3 length | 22.61 (±2.13) | 22.56 | 18.44 | 26.77 |  | 24.36 (±3.98) | 23.89 | 16.55 | 32.17 |
| Thoracic aorta centerline length | 31.78 (±3.01) | 31.70 | 25.87 | 37.69 |  | 35.63 (±5.11) | 35.32 | 25.60 | 45.65 |
| **Curvature** |  |  |  |  |  |  |  |  |  |
| Proximal arch curvature | 0.09 (±0.03) | 0.09 | 0.04 | 0.14 |  | 0.08 (±0.04) | 0.07 | 0.00 | 0.15 |
| Segment 1+2 curvature | 0.09 (±0.03) | 0.09 | 0.04 | 0.14 |  | 0.08 (±0.04) | 0.07 | 0.00 | 0.16 |
| Segment 2+3 curvature | 0.09 (±0.02) | 0.09 | 0.04 | 0.14 |  | 0.07 (±0.04) | 0.07 | 0.00 | 0.15 |
| Thoracic aorta curvature | 0.09 (±0.02) | 0.09 | 0.05 | 0.14 |  | 0.08 (±0.04) | 0.07 | -0.01 | 0.16 |
| **Tortuosity/unfolding** |  |  |  |  |  |  |  |  |  |
| Proximal arch tortuosity | 0.29 (±0.15) | 0.25 | 0.00 | 0.58 |  | 0.32 (±0.13) | 0.29 | 0.06 | 0.58 |
| Arch unfolding | -1.35 (±0.60) | -1.24 | -2.52 | -0.19 |  | -1.80 (±0.59) | -1.73 | -2.96 | -0.65 |
| Segment 2+3 tortuosity | 0.21 (±0.06) | 0.20 | 0.09 | 0.32 |  | 0.19 (±0.07) | 0.18 | 0.07 | 0.32 |
| Thoracic aortic height | 5.17 (±1.16) | 5.20 | 2.91 | 7.44 |  | 6.85 (±1.48) | 6.80 | 3.94 | 9.75 |
| Arch width | 7.46 (±1.29) | 7.39 | 4.94 | 9.99 |  | 7.66 (±1.43) | 7.57 | 4.86 | 10.45 |

Lower stands for lower 95% reference range; Upper stands for upper 95% reference range.
